# Visuomotor interactions in the mouse forebrain mediated by extrastriate cortico-cortical pathways

**DOI:** 10.1101/2022.12.08.519609

**Authors:** Karoline Hovde, Ida V. Rautio, Andrea M. Hegstad, Menno P. Witter, Jonathan R. Whitlock

**Affiliations:** Kavli Institute for Systems Neuroscience, Norwegian University of Science and Technology, NO-7489 Trondheim, Norway; Faculty of Health and Medical Sciences, University of Copenhagen, Blegdamsvej 3B, 2200 Copenhagen N, Denmark

**Keywords:** cortical connectivity, anatomical tracing, visual cortex, extrastriate cortex, frontal cortex, motor cortex

## Abstract

The mammalian visual system can be broadly divided into two functional processing pathways: a dorsal stream supporting visually and spatially guided actions, and a ventral stream enabling object recognition. In rodents, the majority of visual signaling in the dorsal stream is transmitted to frontal motor cortices via extrastriate visual areas surrounding V1, but exactly where and to what extent V1 feeds into motor-projecting visual regions is not well known. To address this we employed a dual labeling strategy in male and female mice in which efferent projections from V1 were labeled anterogradely, and motor-projecting neurons in higher visual areas were labeled with retrogradely traveling adeno-associated virus (rAAV-retro) injected in M2. In flattened sections of dorsal cortex, the most pronounced colocalization V1 output and M2 input occurred in extrastriate areas AM, PM, RL and AL. Coronal sections further showed that neurons in both superficial and deep layers in these regions project to M2, but high resolution volumetric reconstructions revealed that the vast majority of putative synaptic contacts from V1 onto M2-projecting neurons occurred in layer 2/3. These findings support the existence of a dorsal processing stream in the mouse visual system, where visual signals reach motor cortex largely via feedforward projections in anteriorly and medially located extrastriate areas.

**Significance Statement:** Visually guided motor behavior depends on the long-distance relay of signals from visual cortex to frontal motor cortices, but the neuroanatomical connections linking visual and motor systems in rodents are not fully charted. Here, we characterized such pathways by injecting anterograde tracers in primary visual cortex (V1) and retrogradely traveling virus in motor cortex (M2), and visualizing where the projections overlapped. We found preferential colocalization of V1 output and M2-projecting neurons in anteriorly- and medially-located higher visual areas. 3D volumetric reconstructions further showed high rates of putative synaptic connections mainly in superficial layers. Thus, visual signals in the mouse dorsal visual stream reach motor areas at least in part via superficial, feedforward connections in a subset of extrastriate areas.

## Introduction

A common feature of visual cortical organization across mammals is that visual signals from the eye enter primary visual cortex (V1) via the thalamus, then travel to higher visual areas (Benevento & Standage, 1982; Frost & Caviness, 1980; Kaas, 1991; Rosa & Krubitzer, 1999; Salin & Bullier, 1995; Simmons et al., 1982) which process progressively distinct components of visual stimuli (Froudarakis et al., 2019; Maunsell & Newsome, 1987; Nassi & Callaway, 2009; Niell, 2015). In primates and carnivores, higher visual processing streams collect into functionally divergent “dorsal” and “ventral” pathways, with the former supporting spatial and visually guided motor behaviors, and the latter enabling object recognition (Goodale & Milner, 1992; Nassi & Callaway, 2009). Parallel studies in rats, mice and hamsters have also uncovered anatomical (Montero, 1993; Olavarria et al., 1982; Olavarria & Montero, 1989; Wang & Burkhalter, 2007; Wang et al., 2012; Zingg et al., 2014), physiological (Andermann et al., 2011; Garrett et al., 2014; Glickfeld et al., 2013; Han et al., 2022; Marshel et al., 2011; Montero & Jian, 1995) and behavioral (Ho et al., 2011; Kolb & Walkey, 1987; Mlinar & Goodale, 1984; Save et al., 1992; Schneider, 1969; Tees, 1999) evidence supporting a dorsal-*versus*-ventral organization in the rodent visual system, though with fewer functionally specialized nodes. The notion of a dorsal stream in rodents has been further supported by work demonstrating causal contributions of higher visual cortical projections to fine-grained visuomotor control in mice (Itokazu et al., 2018), and has prompted investigations into the role of dorsal stream pathways in the production and perception of naturalistic actions (Tombaz et al., 2020; Viaro et al., 2021) and spatial navigation (McNaughton et al., 1989).

Although in mice it has been established that several of the ∼10 (Froudarakis et al., 2019; Wang & Burkhalter, 2007) higher-visual, or extrastriate, areas send direct projections to frontal and midline motor cortices (Itokazu et al., 2018; Wang et al., 2012; Zingg et al., 2014), several pieces of the puzzle remain missing regarding the anatomical chain by which cortical signaling propagates from V1 to motor areas. For example, it is not known whether the output from V1 is uniformly distributed across frontally-projecting extrastriate cortex, if there are regional preferences among these areas or if there is a laminar profile characteristic of such projections.

We sought to address these questions using a dual pathway tracing approach in which efferent fibers from V1 were labeled using the anterograde tracer 10 KD biotinylated dextran amine (BDA), and motor-projecting extrastriate neurons were labeled via retrogradely traveling, recombinant AAV2/1-retro (rAAV-retro) (Tervo et al., 2016) in secondary motor cortex (M2). Viral injections were targeted to the posterior sector of M2 which receives visual input (Reep et al., 1990; Zingg et al., 2014) and controls movement of the eyes, head and vibrissae (Brecht et al., 2004; Donoghue & Wise, 1982; Sinnamon & Galer, 1984). We visualized the areal overlap of retrograde and anterograde labeling in whole-hemisphere flattened sections of cortex, which revealed a characteristic arrowhead shape of M2-projecting neurons along the anterior perimeter of V1, which overlapped with V1 output fibers in the anteromedial (AM), posteromedial (PM), rostrolateral (RL) and anterolateral (AL) extrastriate areas. In coronal sections, retrograde labeling from M2 appeared columnar in anterior extrastriate areas, spanning superficial and deep layers, and became progressively more superficial at farther posterior locations. Morphological 3D reconstructions revealed substantial putative connectivity between V1 axons and M2-projecting neurons in layers 2/3. Together, our results show that V1 output bound for motor cortex is broadcast non-uniformly across extrastriate regions, and is relayed via feedforward projections largely in layer 2/3.

## Materials and Methods

Eight female C57BL/6JBomTac mice (23-25g, Taconic) and one male C57BL/6JBomTac mouse (33 g) were used in the project. Five animals received injections of virus and tracer into the left hemisphere and their brains were cut in tangential flattened sections. One brain was excluded from the analysis due to poor uptake of the tracer. Four animals received similar injections in the right hemisphere and the brains were cut in coronal sections. One brain was excluded due to a misplaced injection in V1. All animals were housed in single cages, kept on a reversed day-night cycle, and given *ad libitum* access to food and water. All animal procedures were performed in accordance with the Norwegian University of Science and Technology Animal Welfare Committee’s regulations and were approved by the Norwegian Food Safety Authority, and followed the European Communities Council Directive and the Norwegian Animal Welfare Act.

### Retrograde viral tracing and anterograde anatomical tracing

For stereotaxic surgeries, the initial coordinates for V1 and M2 injections were calculated in accordance with Paxinos and Franklin (2012) and adjusted based on previous injections in-house. The animals were deeply anesthetized with isoflurane throughout the surgery and their body temperature was kept stable at 37°C. Local anesthetic Marcain (1-3 mg/kg, bupivacaine, AstraZeneca) was injected above the skull, and analgesics Temgesic (0.1 mg/kg, buprenorphine, Indivior) and Metacam (5 mg/kg, meloxicam, Boehringer Ingelheim Vetmedica) were given subcutaneously. After shaving and disinfecting the head (70% ethanol; iodine, NAF Liniment 2%, Norges Apotekerforening), an incision was made along the midline and the skull was cleaned (hydrogen peroxide, H_2_O_2_; 3%, Norges Apotekerforening), the height of bregma and lambda were measured and adjusted along the anterior-posterior axis to ensure the skull was levelled and two craniotomies were made at the coordinates for injections into secondary motor cortex (M2; AP: +0.3, ML: +0.5, DV: - 0.5) and primary visual cortex (V1; AP: -4.5, ML: +2.3, DV: -0.30-0.60) in either the left (N=5) or right (N=4) hemisphere. A retrograde GFP-tagged adeno-associated virus rAAV2/1-retro (retrograde AAV-CAG-GFP; serotype “retro”, Addgene, Cat. # 37825) was pressure injected into M2 (170, 180, 250, 400 nL volume injections) by use of glass capillaries (World Precision Instruments (WPI), Cat. No. 4878) and Micro4 pump (WPI; speed 35 μL/s), and the capillary was kept in place for 10 min after the injection, to minimize leakage of the virus. An anterograde tracer, 10 KD biotinylated dextran amine (BDA, Dextran, Biotin, 10,000 MW, Lysine Fixable (BDA-10,000), Thermo Fisher Scientific Cat. No. D1956, RRID:AB_2307337 in 5% solution in 0.125 M phosphate buffer), was injected into V1 iontophoretically by pulses of positive DC-current (6 s on/off alterations, 6 μA, 10 min) using glass micropipettes (20 μm tip, Harvard apparatus, 30-0044). After the injection was completed, the craniotomies were filled with Venus Diamond Flow (Kulzer, Mitsui chemical group, Cat. # 879566), the skull was cleaned and the skin was sutured and disinfected with iodine. The animal was kept in a heated chamber until awake and active. Post-operative analgesic (Metacam; 5 mg/kg) was given 12 hours post-surgery and the health of the animal was closely monitored the days after surgery.

### Perfusion and tissue processing

All animals were killed and perfused 21 days post-surgeries.

#### Tangential flattened sections

The animals that received injections in the left hemisphere were given an overdose of pentobarbital (0.2 mL / 100 g) and transcardially perfused using fresh ringer’s solution (0.025% KCl, 0.85% NaCl, 0.02% NaHCO_3_, pH 6.9) and PFA (1%, 0.125 M phosphate buffer, pH 7.4), and the brains were carefully removed and kept in a cup of PFA. Within one hour, the cortex of the left hemisphere was dissected out and flattened, and tangential sections (50 μm) were prepared. To do so, the intact brain was cut along the midline, subcortical areas and cerebellum were removed, and one cut was made in the fornix dorsal to the anterior commissure. Horizontal cuts were then made along the white matter, and relief cuts were made ventral to postrhinal cortex and in the anterior cingulate cortex. The hippocampus was unfolded, and the cortex was flattened between two microscope glasses covered with parafilm (Laboratory film, Pechiney, Plastic packaging, Chicago) and submerged in PFA (4%) overnight at 4°C with a glass weight on top (52 g). The following day, the flattened cortex was removed from the microscope slides and left in a cryoprotective dimethyl sulfoxide solution (2% dimethyl sulfoxide, DMSO, in 0.125 M phosphate buffer; VWR) overnight. The flattened cortex was then cut in 50 μm tangential sections in one series on a freezing microtome (Microm HM430, Thermo Scientific, Waltham, USA).

#### Coronal sections

Following the same procedure as above, animals with right hemisphere injections were perfused with fresh ringer’s solution and PFA (4%). The brain was placed in a container with PFA (4%) overnight, transferred to cryoprotective solution (DMSO, 2%) and stored overnight. The brain was cut on a freezing microtome in 40 μm sections in three series. The first series was mounted on Superfrost Plus microscope slides (Gergard Menzel GmbH, Braunschweig, Germany) and used for Nissl staining, the second was processed to reveal the tracer and virus, and the third was stained with 3.3′-Diaminobenzidine tetrahydrochloride (DAB, Sigma-Aldrich, St. Louis, USA) against the muscarinic acetyl choline receptor 2 (M2AChR2) or kept as a backup in cryoprotective solution stored at −24°C.

### Histology and immunohistochemistry

*Nissl*. Series one of the coronal sections was stained with Nissl staining. To do so, sections were hydrated in running water and dehydrated in baths with increasing percentage of ethanol (50%, 70%, 80%, 90%, 100% x3), cleared in a solution of xylene (2 min; VWR, International, Fontenay-sous-Bois, France) and rehydrated in decreasing percentage of ethanol, followed by a brief rinse in running water prior to staining in Cresyl violet on a shaker (3 min). The sections were rinsed in water, differentiated in a solution of ethanol/acetic acid (0.5% acetic acid in 70% ethanol; VWR, International, Fontenay-sous-Bois, France) until reaching the desired staining contrast, and cleared in two xylene baths (2 min, 20 min) before being coverslipped with an entellan-xylene solution (Merck KGaA, Darmstadr, Germany).

#### BDA visualization and enhancement of rAAV-retro signal

All flattened tangential sections and series two of the coronal sections were processed to reveal the BDA tracer and to enhance signal from the virus using a two-day immunohistochemical procedure. On day one, the sections were washed in phosphate buffered saline (PBS; 3 × 5 min), followed by a phosphate buffered saline solution with Triton (PBS 0.1M, 0.3% Triton, 3% BSA; 2 × 10 min) on a shaker (100 rpm) at room temperature (RT). The sections were incubated with anti-GFP primary antibody (GFP; rabbit anti-GFP, 1:1000, ThermoFisher Scientific, A-11122) overnight on a shaker (60 rpm) at 4°C. On day two, the sections were washed in PBS solution (PBS 0.1M, 0.3% Triton, 3% BSA; 2 × 5 min) and incubated with secondary antibody (AlexaFluor 488-tagged goat anti-rabbit Ab, 1:1000, ThermoFisher Scientific, A-11008) and with Alexa Fluor 633-conjugated Streptavidin (1:400, Thermo Fisher Scientific Cat. No. S-21375, RRID:AB_2313500) against BDA on a shaker (60 rpm) at RT (75 min). The sections were rinsed in Tris buffer 0.606% (Tris(hydroxymethyl)aminomethane, pH 7.6; 3 × 10 min), mounted on non-frost microscope slides using a Tris-gelatin solution (0.2% gelatin in Tris-buffer, pH 7.6) and coverslipped with an entellan-xylene solution.

#### DAB staining against M2AChR

Series three of the coronal sections were stained with 3.3′-Diaminobenzidine tetrahydrochloride (DAB, Sigma-Aldrich, St. Louis, USA) to visualize M2AChR density and were used only for delineation purposes. To do so, sections were rinsed in PBS (0.125 M, 2 × 5min) followed by TBS-Tx (2 × 5 min) and incubated with primary antibody (Rat anti-muscarinic M2 monoclonal antibody, unconjugated, clone m2-2-b3, 1:750, Millipore Cat. No. MAB367, RRID:AB_94952; overnight at RT), washed in TBS-Tx (2 × 5 min) and incubated with mouse-absorbed, rabbit-anti-rat secondary antibody (Anti-rat IgG (H+L), 1:300, Vector Laboratories Cat. No. BA-4001, RRID:AB_10015300) for 90 min at RT. The sections were washed in TBS-Tx (2 × 5 min), in PB (2 × 5 min), in H_2_O_2_-metanol solution (0.08%, Sigma-Aldrich, 2 × 5 min) and in TBS-Tx (2 × 5min) and incubated with a Vector ABC kit (Vector laboratories, Inc., Burlingame, USA) for 90 min at RT, per the manufacturer’s instructions. They were then washed in TBS-Tx (2 × 5 min) and Tris-buffer (2 × 5 min) before being incubated with DAB (10 mg in 15 mL Tris-buffer, Sigma-Aldrich) at RT. H_2_O_2_ (2 μL, 30%, Sigma-Aldrich) was added to the DAB solution immediately prior to the incubation. The solution was filtered, the sections were incubated in DAB until the desired level of staining was reached and washed in Tris-buffer solution. A 0.2% gelatin solution was used to mount the sections on Menzel glass slides, the slides were dried overnight on a heated pad and coverslipped with an entellan-xylene solution.

### Imaging and analyses

All Nissl and M2AChR stained sections were digitized for analyses using a bright field scanner (Zeiss Axio Scan.Z1). Sections with fluorescence labeling were examined in a fluorescence microscope (Zeiss Axiomager M2) and digitized with a fluorescence scanner (Zeiss Axio Scan.Z1). Lower exposure time was used for the sections with the injection sites in V1 and M2 to avoid saturation of the signal.

High resolution images (63x oil) in z-stacks (typically 70-90 planes, 0.14μm intervals, 0.05μm pixel size) were taken of selected sections with fluorescence labeling using a Zeiss confocal microscope (LSM800). The images we deconvoluted in Huygens 19.10 (Scientific Volume Imaging) using the default express deconvolution. The deconvoluted image stack was saved as 16 bit .pic files (one for each fluorescent channel) and opened in Neurolucida360 (MBF Bioscience) for reconstruction.

The outlines of V1 and S1B were drawn on flattened tangential sections using myeloarchitectonic features visible in layer IV (Extended Data Fig. 1-1). The same outlines were copied and overlaid on sections cut through superficial layers of cortex (Fig.1), where myeloarchitectonic features were not present.

**Figure 1.**
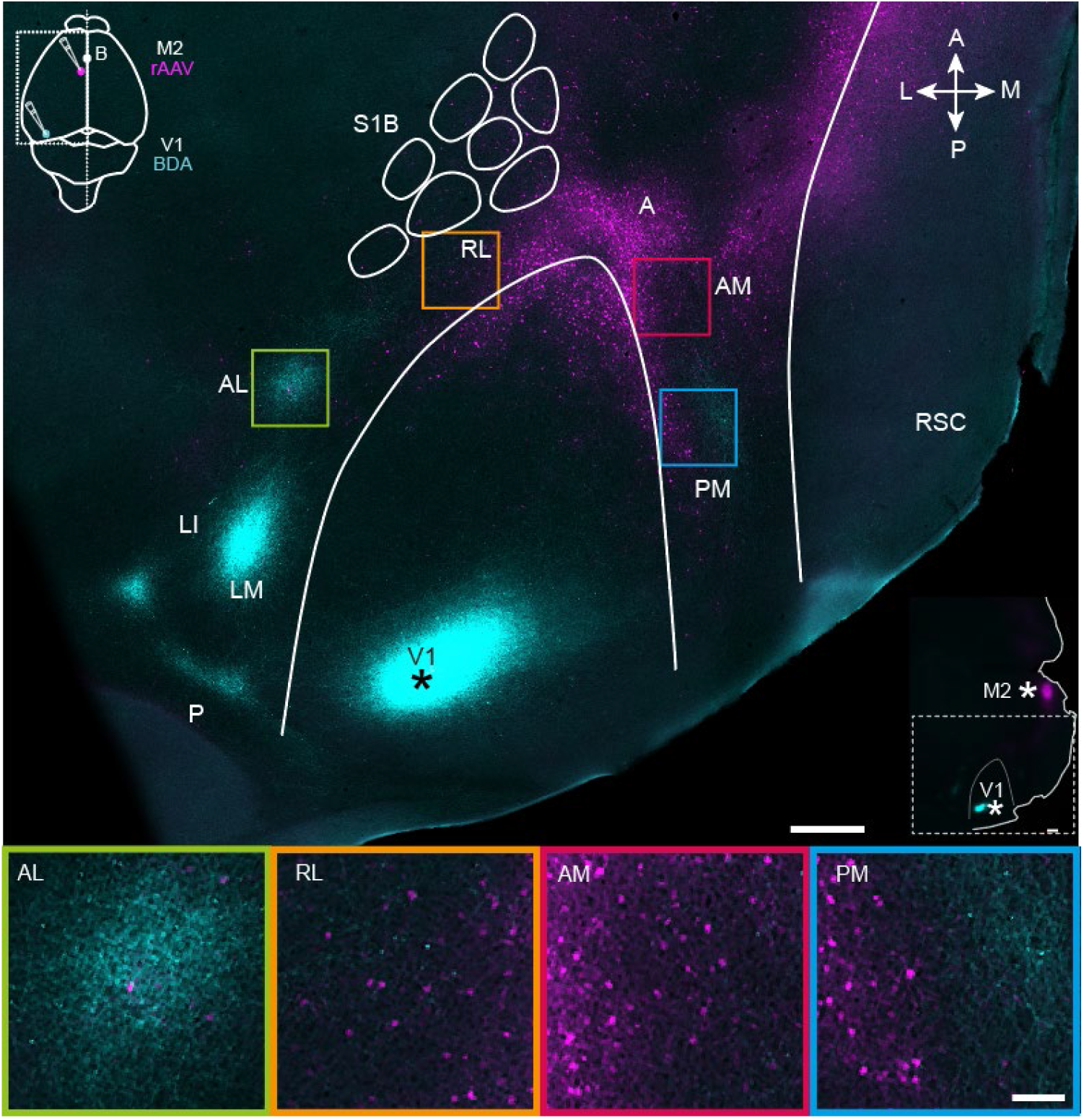
Projections from V1 (cyan) and M2-projecting neurons (magenta) viewed in a representative tangential section through superficial layers in flattened dorsal cortex. Top, A tangential section through layer 2/3 shows the BDA injection site in V1 (marked with asterisk; see also inset at top left) and projections to extrastriate areas at the periphery of V1. Projection neurons targeting M2 are shown in magenta (injection schematic shown in inset), and colocalized with V1 efferent fibers in areas AL, RL, AM and PM. The outlines of V1 and the S1 barrel fields were traced using myeloarchitectonic patterns and M2AChR staining from a neighboring section in layer 4 (Methods). Note, a shorter exposure time was applied when scanning the injection site than for projections to avoid signal saturation (Methods). Bottom, magnification of extrastriate areas highlighted in the flattened section above. See list for abbreviations. White scale bar in upper image = 500 μm; in lower image = 100 μm.

### Reconstruction and proximity testing

The deconvoluted image stacks obtained from Huygens 19.10 (see above) were opened and the two fluorescence channels were merged in Neurolucida360. The black point of the image was increased 10%, the white point lowered 90% and gamma was set to 1.20 for visualization purposes, as this enhanced image contrast and removed background noise. Dendrites were traced using the “user-guided tracing” mode with the method “voxel scooping” before spines were detected using the nearest branch mode. Next, axons were traced using the tracing option “direction kernels” and boutons were detected using the nearest branch mode. After the reconstruction was complete, synaptic markers were placed using the “synaptic markers” button with a 0.25μm requirement and the results were saved as a .DAT file. The file was opened in Neurolucida explorer and a branch structure analysis was performed using the synapses mode and synaptic markers details. The markers with a distance below 0.25μm were considered as putative synaptic contacts and used in subsequent comparisons between areas. The soma was not reconstructed in Neurolucida but imported as a 2D image into the final 3D reconstruction of neurons for illustration purposes only.

## Results

### Areal organization of V1 output and M2 input

First, we sought to gain an overview of cortical regions where efferent fibers from V1 and cell bodies of motor-projecting extrastriate neurons were colocalized. To do so, four mice received unilateral injections of BDA targeted to the posterior pole of V1 (Fig. 1, top panel), which previous work has shown sends projections to all downstream visual areas (Olavarria & Montero, 1989; Wang & Burkhalter, 2007). The same animals received rAAV-retro injections in the posterior sector of secondary motor cortex, M2 (per the nomenclature of (Paxinos, 2012)), due to the high density of visual input it receives in mice and rats (Reep et al., 1990; Wang et al., 2012). Three weeks after surgery, the brains were removed and the left hemisphere was dissected out, flattened and cut tangentially into sections parallel to the brain surface, allowing us to visualize regional labeling of efferent V1 fibers and M2-projecting neurons in extrastriate cortices. At least seven extrastriate areas were discernable based on the topographical positioning and orientation of projection plexuses relative to V1, with the most prominent labeling from V1 in LM, LI, AL and PM, with more moderate labeling in RL, A and AM (Fig. 1, Top; regional nomenclature per (Olavarria et al., 1982) and (Wang & Burkhalter, 2007)). The location of V1 projections to extrastriate areas was consistent across mice, though the relative strength of labeling within regions varied depending on the mediolateral location of the injections in V1 (Extended Data Fig. 1-1) and was therefore not quantified here, but can be found in earlier work (Wang & Burkhalter, 2007).

Retrogradely labeled M2-projecting neurons were condensed at the anterior pole of V1, and flanked its medial and lateral borders in a V-shape that could continue as far laterally as area AL, and as far posteriorly as area PM (Fig. 1, top; Extended Data Figs. 1-1 and 1-2). Regions showing the most extensive coincident labeling from V1 and to M2 were AM, PM and RL, which partly overlapped with posterior parietal cortex (PPC) (Gilissen et al., 2021; Hovde et al., 2018), and sparse labeling of M2-projecting neurons was present area AL. Labeling in area A was weak, which could have been due to the locations of our injections in V1 (Wang & Burkhalter, 2007). But in view that this was observed consistently across animals, we find it likely that overlap and possible contacts between V1 axons and M2 projection neurons were generally rare in area A. In all extrastriate regions, dual labeling of V1 efferent fibers and M2-projecting neurons was strongest in superficial layers and layer 4, with sparser labeling in deeper layers (Extended Data Fig. 1-2); this pattern was investigated in more detail in coronal sections.

### Laminar organization of V1 output fibers and M2-projecting neurons

Similar injections of BDA and rAAV-retro were made in the right hemisphere of V1 and M2 in four additional mice, and coronal sections were collected in cortical regions spanning anteriorly from M2 to the posterior extent of V1 (Fig. 2, right, Extended Data Figs. 2-1 and 2-2). Consistent with our observations in flattened sections, regions with the densest axonal plexuses from V1 were LM, LI, AL and PM, with more moderate but clear labeling in AM and RL (Fig. 2 left, Extended Data Figs. 2-1 and 2-2). V1 projections were observed in both superficial and deep layers of all extrastriate areas, though across animals we noted axonal fibers were concentrated in layers 3 and 5 (Fig. 2, left, magnifications; Extended Data Figs. 2-1 and 2-2). As with flattened sections, there were very few axonal fibers from V1 in area A.

**Figure 2.**
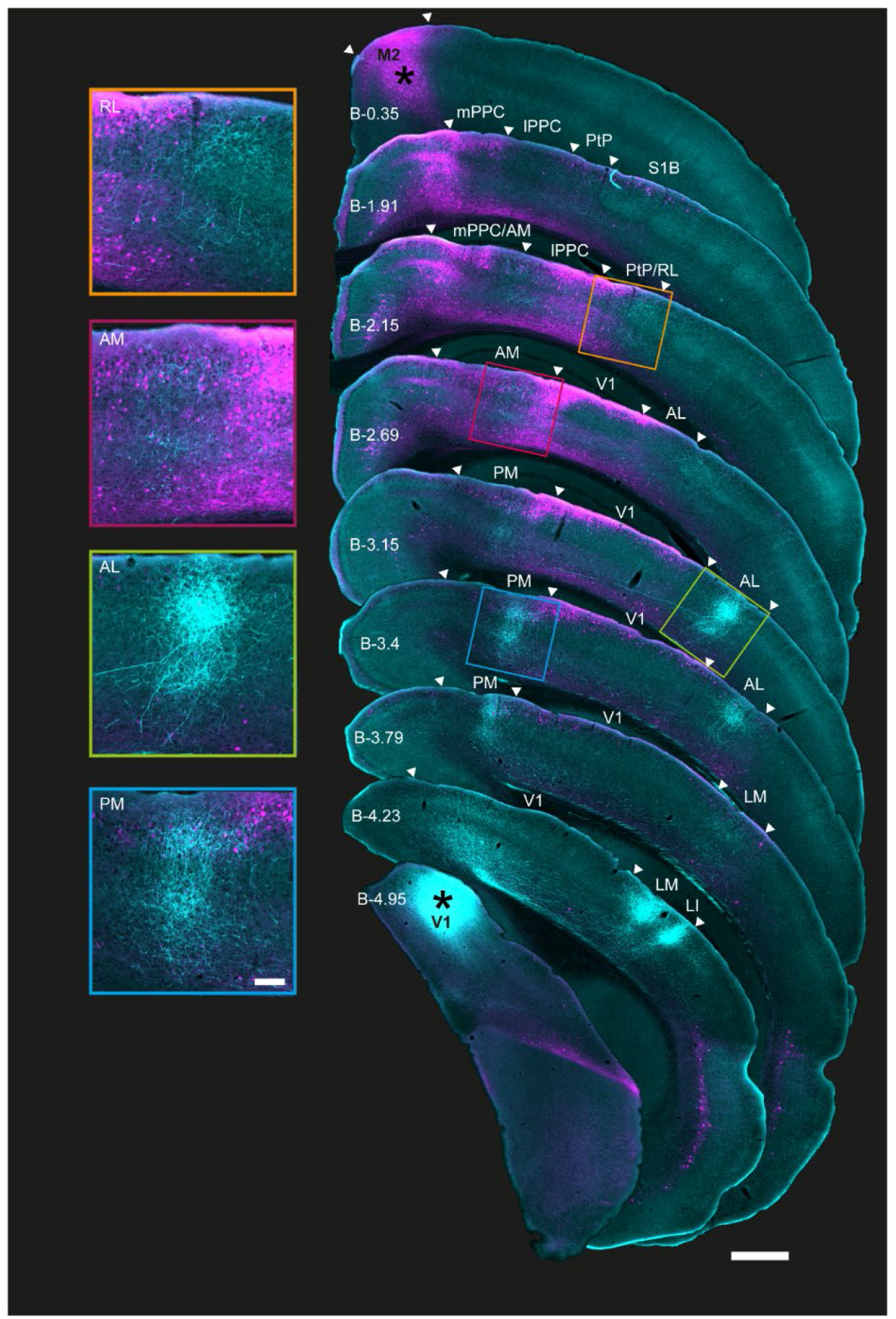
Coronal sections showing the laminar profile of BDA-labeled V1 projections and rAAV-retro-labeled M2-projecting neurons in posterior cortices. Right, low magnification series of coronal sections from the right hemisphere arranged from anterior (top) to posterior (bottom); injection sites marked with asterisks. Extrastriate areas and PPC boundaries are indicated by white triangles; PPC, its sub-areas, and V1 were delineated using adjacent Nissl and immunohistochemically stained sections from the same series. Anterograde and retrograde labeling from V1 and M2 colocalized in areas RL, AM and PM, including in PPC, as well as area AL. M2 projecting neurons were found in superficial layers in all regions in the series, and extended to deep layers at more anterior locations, especially in RL and AM. Left, magnified view of extrastriate areas highlighted in sections to the right. As with Figure 1, shorter exposure times were used for injection sites than projections to avoid signal saturation (Methods); the figure is for illustration purposes. See list for abbreviations. Approximate bregma coordinates (B; Paxinos and Franklin, 2012) are noted on each section; scale bar = 500μm.

Across animals, rAAV-retro labeling from M2 was abundant in several extrastriate regions and formed apparent columns spanning from layers 1 to 6 in more anterior regions. At its anterior extent, retrograde M2 labeling formed a column at the border of the medial PPC (mPPC) and agranular retrosplenial cortex (RSA; Fig. 2, 2^nd^ coronal section), then, progressing posteriorly, split into medial and lateral branches that overlapped with mPPC and AM medially, and lateral PPC (lPPC) and RL laterally. Retrograde labeling from M2 was strong in superficial and deep layers in all PPC sub-regions as well as AM and RL as far posterior as Bregma (B) level -3.15 (Fig 2.; Extended Data Figs. 2-1 and 2-2, 4^th^ and 5^th^ coronal sections; Bregma location based on (Paxinos, 2012)). Further posteriorly, strong retrograde labeling was observed mostly in superficial layers 1-3 (≥ B -3.15) in PM. Laterally, a similar pattern was observed in AL and V1, with the exception that layer 5 was almost devoid of neuronal labeling and layer 6 only showed sparse neuronal labeling (Fig. 2, 5^th^ and 6^th^ sections; magnified insets, left). Retrograde labeling tapered off completely in all layers at the farthest posterior locations, with no M2-projecting neurons remaining in areas LM or LI, or at the posterior extent of V1 (≥ B -4.23; Fig. 2, right). Thus, of the six identifiable extrastriate areas in this tissue series, regions RL, AM, PM, and AL, in addition to the anterior-most portion of V1, were potential nodes at which visual signals were conveyed to secondary motor cortex.

### Neuronal reconstructions and localization of putative synaptic contacts

The colocalization of V1 fibers and M2-projecting neurons in specific extrastriate areas suggested they could serve as nodes for the propagation of visual information to the motor system, but overlap *per se* does not mean that synaptic connections are in fact present. To test this possibility, we stacked high-magnification (63x) serial confocal scans and created 3-dimensional morphological reconstructions in all extrastriate areas in which M2-projecting neurons had clearly evident pre-synaptic V1 fibers in their vicinity (Fig. 3A; Methods). We note that the resulting reconstructed cells were biased in that they were chosen from areas that appeared to have colocalized labeling. Neurons were selected for reconstruction based on three criteria: (i) the presence of a clear and completely filled neuronal soma associated with (ii) filled, long spiny dendrites, and (iii) a sufficiently densely labeled axonal plexus from V1 overlapping the dendrites, where connectivity was expected to occur (see examples in magnified insets in Figure 3B). Putative synapses were identified using close spatial proximity (< 0.25μm) between axonal varicosities and dendritic spines as a proxy for synaptic contacts ((Methods); (Koganezawa et al., 2015; Wouterlood et al., 2008)), and group data were generated for areas AM (n = 8 neurons from 3 mice), AL (7 neurons, 2 mice), PM (7 neurons, 3 mice).

**Figure 3.**
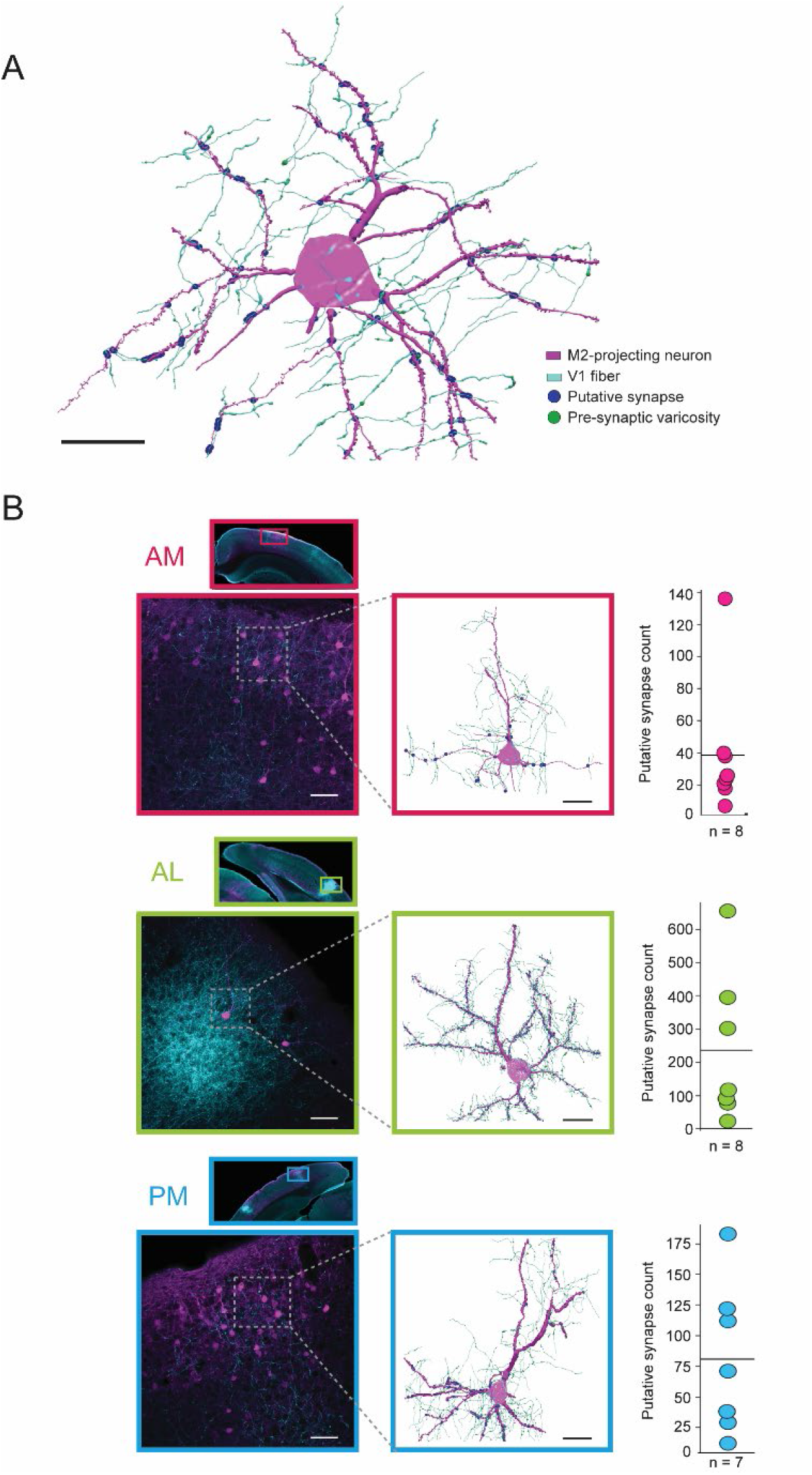
Anatomical reconstructions and proximity analysis of V1 axons and M2-projecting neurons in extrastriate areas AM, AL and PM. A. Representative reconstruction of a single M2-projecing pyramidal neuron from area AL (magenta) receiving synaptic input from V1 axons (cyan); putative synaptic contacts are shown as blue circles and pre-synaptic varicosities are visualized as green enlargements on the V1 axons. They are not visible at points of putative contact due to the overlapping blue circles. Black scale bar = 20 μm. **B**. Top row, inset, a low magnification scan of a coronal tissue section highlighting the portion of area AM where the reconstruction was performed. Top row, lower left, a higher magnification field of view of area AM shown above. Top row, middle, reconstruction of the neuron in the stippled box to the left, with the same labeling convention as the example in **A**. Top row, right, mean (solid line) and distribution of putative synapses for all reconstructed neurons in area AM. Middle row, same as top, but for area AL. Bottom row, same upper rows, but for area PM. Black scale bar = 20 μm. White scale bars = 50 μm.

Area AM, which had high density retrograde labeling from M2, had the lowest mean number of putative V1 synapses per neuron (Fig. 3*B*, top; mean = 38.8 ± 13.7 SEM; median = 25; range of 7 to 136 per neuron; all reconstructed AM neurons shown in Extended Data Fig. 3-1). Area AL, on the other hand, typically contained few M2-projecting neurons, but had the highest rate of V1 contacts of the 3 regions (Fig. 3*B*, middle; mean = 237.1 ± 89.6; median = 117; range of 23 to 655 per neuron; all remaining reconstructions in Extended Data Fig. 3-2). In PM, the number of synaptic connections from V1 were intermediate between AM and AL across animals (Fig. 3*B*, bottom; mean = 81 ± 28.6; median = 71; range of 12 to 183; Extended Data Fig. 3-3). We also note that, across all regions and animals, the large majority of neurons were reconstructed from superficial layers (n = 6 of 8 neurons in AM; 6 of 7 in AL; 6 of 7 in PM), since retrograde M2 labeling was characteristically sparser in deeper layers, and very few incidents of nearby axonal processes from V1 in layers 5 and 6 (Extended Data Fig. 3-1, 3-2 and 3-3).

## Discussion

We used a dual anterograde and retrograde labeling strategy to characterize the regional intersection and connectivity patterns of V1 output fibers onto motor cortical-projecting neurons in mouse extrastriate cortex. We prepared flattened sections, which provided an overview of the entire dorsal cortex, and found that visual and motor pathways overlapped specifically in extrastriate and posterior parietal regions which support visuospatial and motor behavior in rodents (L. L. Chen et al., 1994; Harvey et al., 2012; Itokazu et al., 2018; Kolb et al., 1994; Kolb & Walkey, 1987; McNaughton et al., 1989; Nitz, 2006; Save et al., 1992; Schneider, 1969; Tees, 1999; Whitlock et al., 2012; Wilber et al., 2014; Yang et al., 2017). Co-labeling in these regions was confirmed in coronal sections, which revealed a columnar-like organization of M2-projecting neurons across deep and superficial layers in AM and RL which receded into mainly superficial layers posteriorly in PM, AL and in V1. A higher resolution analysis of reconstructed neurons further showed that the majority of identifiable putative synaptic connections between V1 and M2-projecting neurons occurred in layer 2/3.

Though our observations were consistent across animals, there are methodological limitations in the study which could influence the patterns of connectivity we observed. In visual cortex, for example, the posterior location of the injection produced strong labeling in LM, LI and AL, whereas weak labeling was observed in area A, and the medial-lateral location of the injections likely shaped the spatial distribution of labeling within extrastriate areas (Extended Data Figs. 1-1, 2-1, 2-2) (Hovde et al., 2018; Wang & Burkhalter, 2007). The retrograde labeling from M2 was also restricted by the use of single injections, which were targeted to the posterior sector due to its known connectivity with visual regions (Reep et al., 1990; Zingg et al., 2014), though this could have biased the distribution of labeling toward medial extrastriate and PPC subregions (Olsen et al., 2019). The injections themselves in V1 and M2 were also densest in intermediate layers (mainly layers 2-5; Extended Data Figs. 2-1 and 2-2), which would favor labeling in matching layers in either up- or down-stream regions (Callaway, 2004; Felleman & Van Essen, 1991). Thus, although we directly observed robust feedforward projections from V1 to extrastriate layers 2/3, projections from V1 to the tips of apical dendrites of layer 5 neurons in layer 1 may have been underrepresented. Because of these constraints, the present study was intended to provide a description of the patterns of coincident anterograde and retrograde labeling, rather than a quantitative account of projection densities or the directionality of connections between visual and motor areas.

Nevertheless, the conjoint use of anterograde and retrograde labeling afforded a direct snapshot of the anatomical organization of primary visual outputs onto motor cortical inputs, which occurred in the same extrastriate regions hypothesized to comprise the dorsal visual processing stream in rodents (Wang et al., 2011; Wang et al., 2012).

Intratelencephalic (IT)-to-IT projections, which originate mainly in layers 2/3 and layer 5 (Douglas & Martin, 2004; Harris & Shepherd, 2015), were labeled strongly in our preparations, and the presence of M2-projecting neurons in layers 2/3 and layer 5 suggests that extrastriate and M2 regions participate in mutual feed-forward and feed-back projection pathways (Callaway, 2004; Douglas & Martin, 2004), putting them at nearby stages in the cortical processing hierarchy (Felleman & Van Essen, 1991; Hilgetag et al., 1996). Indeed, previous work using dual anterograde tracers showed that regions RL and AM are reciprocally connected with M2, and that all three regions shared directional preferences for visual stimuli and eye movements, suggesting that they comprise an extended sensory-motor network (Itokazu et al., 2018). The frontally projecting neurons we observed in superficial V1 could also participate in mutual feedback loops with frontal motor areas. Based on existing work, such anatomical loops appear to support visual attention (Zhang et al., 2014) as well as predicting expected changes in visual flow due to self-generated movement (Keller et al., 2012; Leinweber et al., 2017), though for now the exact contributions of these projections are yet to be demonstrated.

Although our results provide support for the existence of specialized visual processing streams in rodents, the degree of correspondence with monkeys and carnivores is limited due to considerable differences in brain size, interconnectivity and the relative simplicity of cortical hierarchies in rodents compared to mammals with more differentiated cortices (Coogan & Burkhalter, 1993; Felleman & Van Essen, 1991; Krubitzer & Seelke, 2012; Kaas, 1991). For example, whereas visual cortex in primates projects only to nearby higher visual areas and area MT (Felleman & Van Essen, 1991; Hilgetag et al., 2000), V1 in rodents projects to a host of non-visual regions including somatosensory, cingulate, retrosplenial and postrhinal cortices (Burwell & Amaral, 1998; Miller & Vogt, 1984; van Groen & Wyss, 1992; Vogt & Miller, 1983), suggesting a wider and more direct intermixing of visual signals across sensory and cognitive modalities. Thus, visuomotor (Rozzi et al., 2008) and spatial functions (Chafee & Crowe, 2012; Crowe et al., 2005), as well as the reference frames in which they are encoded (X. Chen et al., 2013), are likely follow tighter and more organized anatomical localization in primates and carnivores which, in rodents, appear distributed over coarser topographies. Our present results nevertheless confirm that the anterior and medial extrastriate areas, in at least the superficial layers, are key sites of visuomotor integration which like subserve a variety of visually- and spatially guided behaviors.

## Acknowledgements

We thank M. Andresen, G. M. Olsen, P. Girao and K. Moan for technical assistance; and M.J. Nigro for helpful discussions and comments on the manuscript.

## Extended data figures

**Extended Data Figure 1-1.**
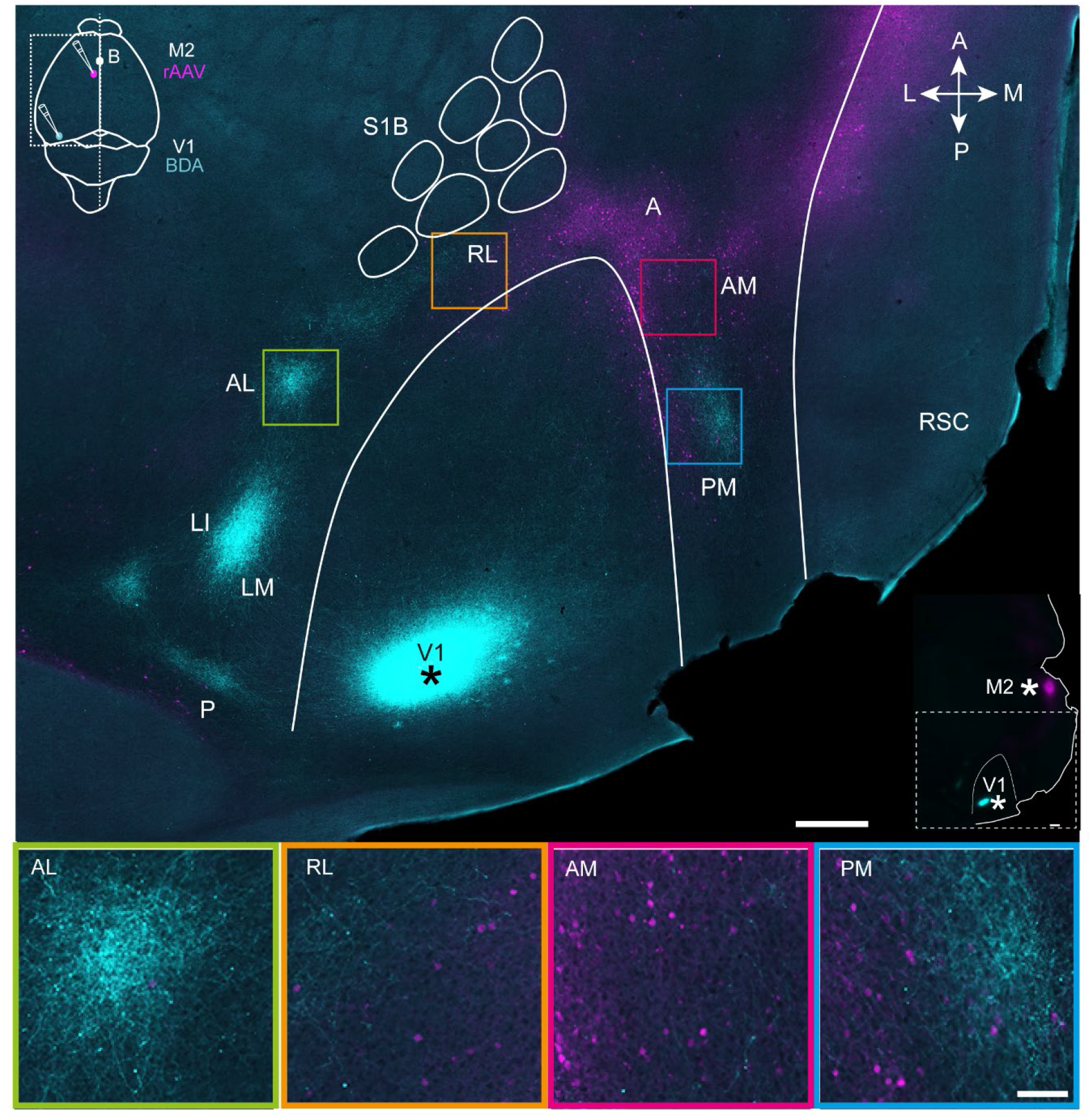
Projections from V1 (cyan) and M2-projecting neurons (magenta) viewed in a tangential section of flattened dorsal cortex of same brain as Figure 1 (mouse #83487). Top, a tangential section through layer 4 of flattened cortex shows the BDA injection site in V1 (marked with asterisk; see also inset at top left) and projections to extrastriate areas at the periphery of V1. M2-projecting neurons are shown in magenta (see injection schematic in inset), and colocalized with V1 efferent fibers in areas AL, RL, AM and PM. The outlines of V1 and the S1 barrel fields were traced using myeloarchitectonic patterns and M2AChR staining (Methods) and copied over to the neighboring section in Figure 1. When scanning the section, shorter exposure time was applied for the injection site than for projections to avoid saturation of signal (Methods). Bottom, magnification of extrastriate areas highlighted in the flattened section above. See list for abbreviations. White scale bar in upper image = 500 μm; in lower image = 100 μm.

**Extended Data Figure 1-2.**
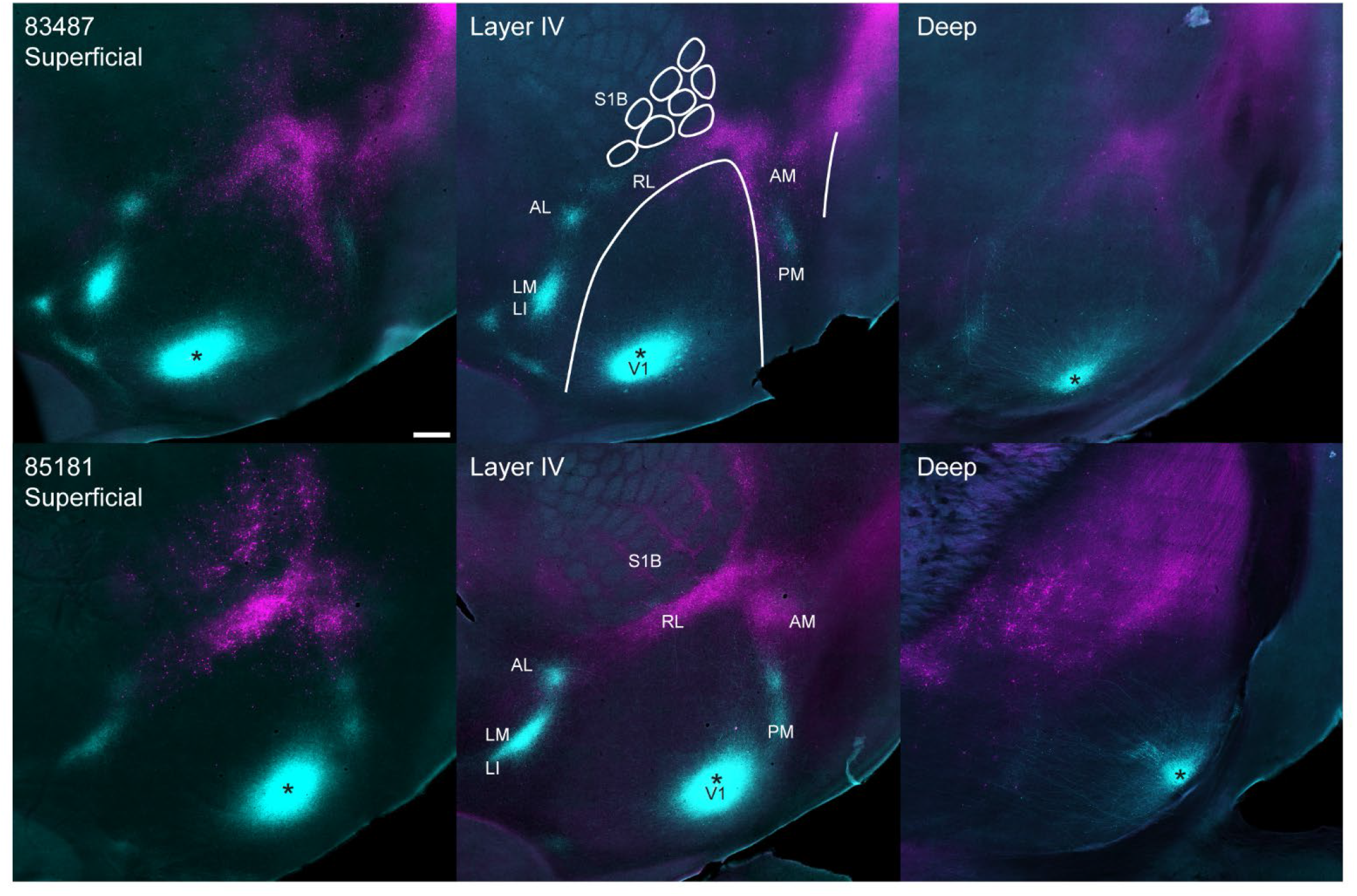
Projections from V1 (cyan) and M2-projecting neurons (magenta) viewed in three dorsoventral levels of tangential sections through flattened dorsal cortex. Top (left to right), superficial layers, layer 4 and deep layers of the same brain from Figure 1 (mouse #83487). Outlines of V1 and S1B were drawn using myeloarchitectonic patterns from layer 4 (top middle); the injection site in V1 is marked with asterisk. Bottom (left to right), superficial layers, layer 4 and deep layers from a second mouse brain (#85181) with a more medial injection site in V1, indicated with asterisk. See list for abbreviations. Scale bar = 500 μm.

**Extended Data Figure 2-1.**
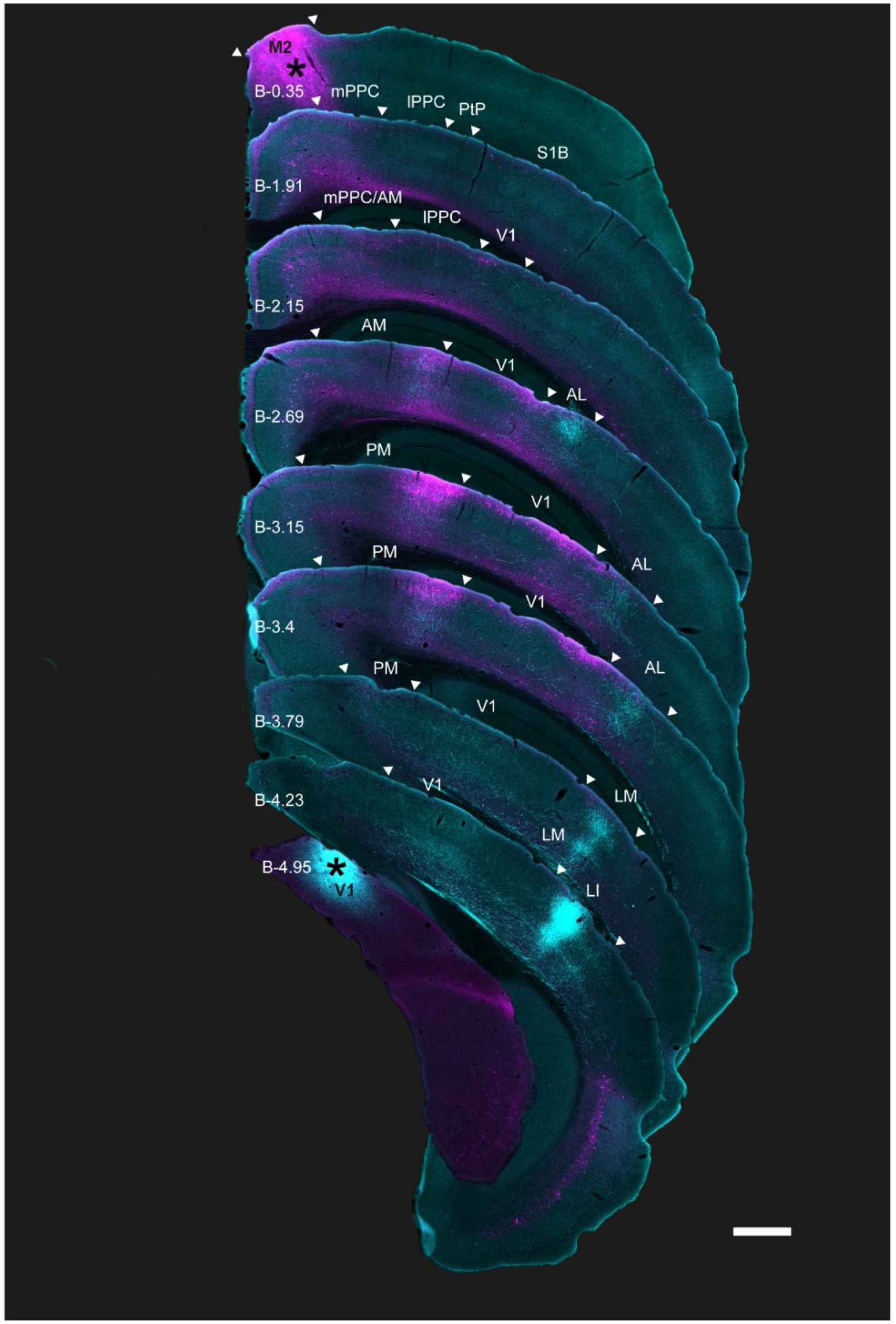
Coronal sections showing the laminar profile of BDA-labeled V1 projections and rAAV-retro-labeled M2-projecting neurons in posterior cortices (mouse #81236). A low magnification series of coronal sections arranged from anterior (top) to posterior (bottom) from the right hemisphere; injection sites in M2 and V1 are marked with asterisks. Extrastriate and PPC boundaries are indicated by white triangles; PPC, its sub-areas, and V1 were delineated using adjacent Nissl and immunohistochemically stained sections from the same series (Methods). As with Figure 1, shorter exposure times were used for injection sites than for projections to avoid signal saturation (Methods); the figure is for illustration purposes. See list for abbreviations. Approximate Bregma coordinates (per Paxinos and Franklin, 2012) are noted on each section; scale bar = 500μm.

**Extended Data Figure 2-2.**
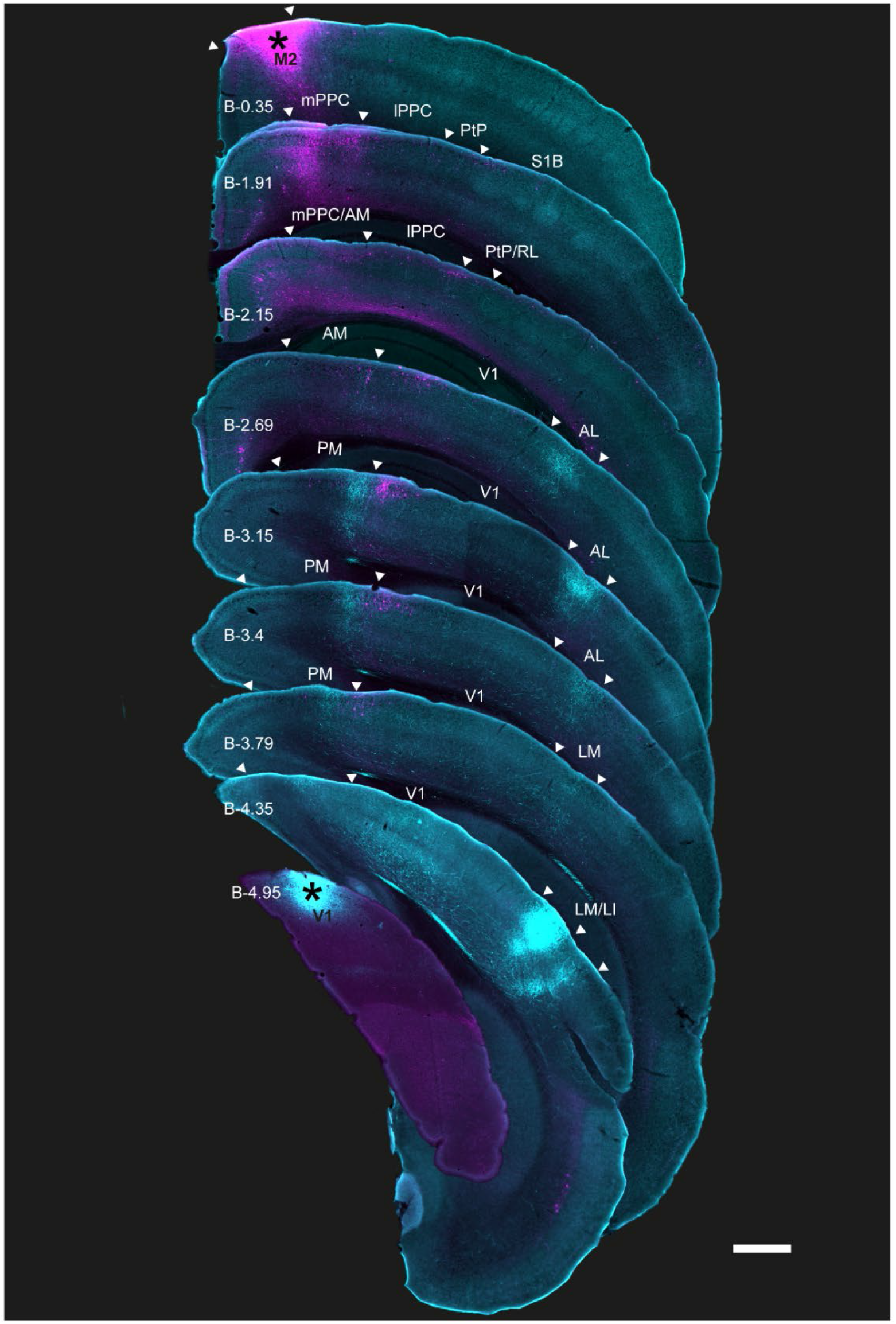
Coronal sections showing the laminar profile of BDA-labeled V1 projections and rAAV-retro-labeled M2-projecting neurons in posterior cortices (mouse #81233). Low magnification series of coronal sections arranged from anterior (top) to posterior (bottom) from the right hemisphere; injection sites marked with asterisks. Extrastriate and PPC boundaries are indicated by white triangles; PPC, its sub-areas, and V1 were delineated using adjacent Nissl and immunohistochemically stained sections from the same series (Methods). As with Figure 1 and Extended Data Figure 2-1, shorter exposure times were used for injection sites than for projections to avoid signal saturation (Methods); the figure is for illustration purposes. See list for abbreviations. Approximate Bregma coordinates (Paxinos and Franklin, 2012) are noted on each section; scale bar = 500μm.

**Extended Data Figure 3-1.**
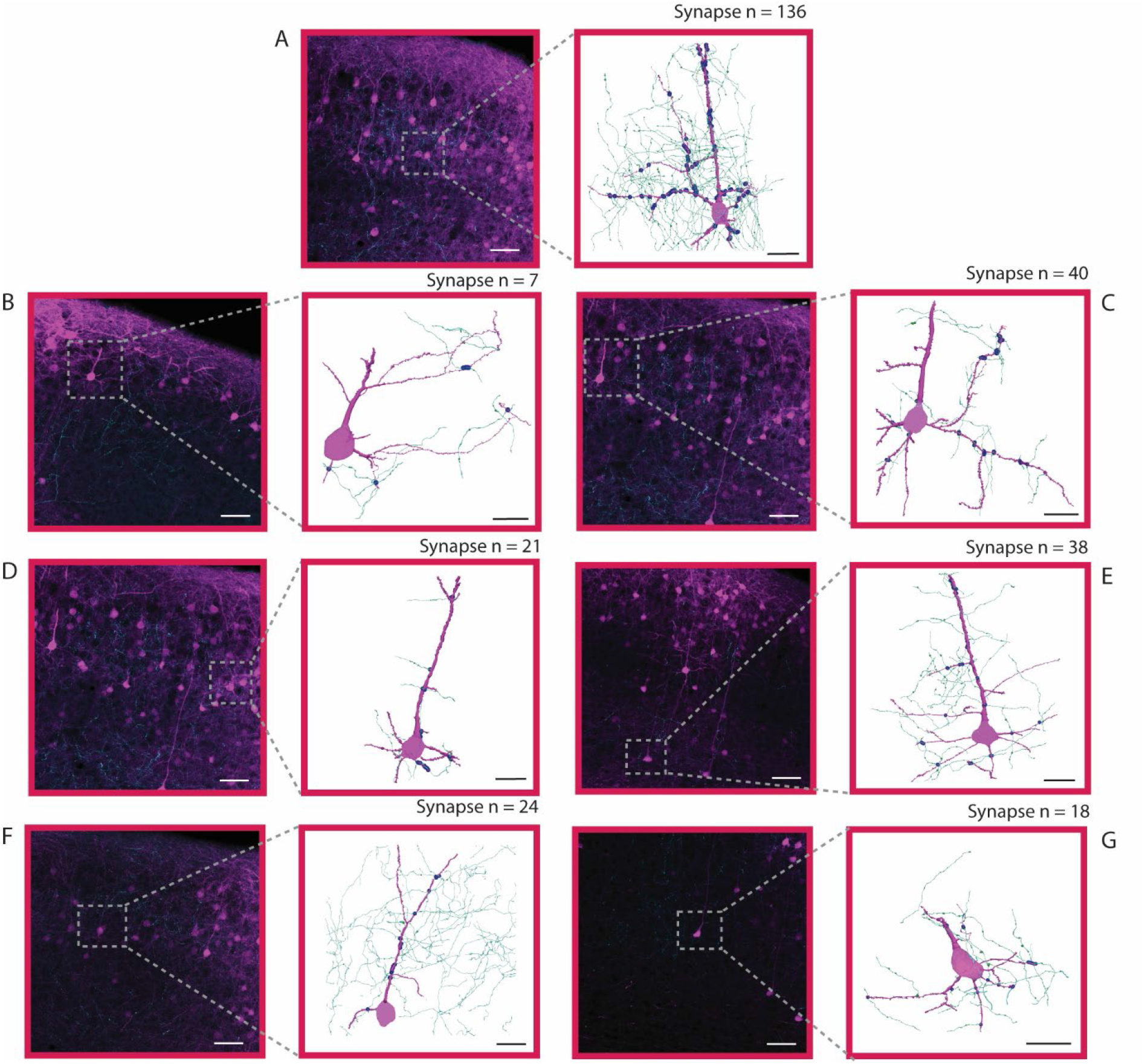
Anatomical 3D reconstructions of V1 axons and seven M2-projecting neurons in extra striate area AM. Left, inset, 20x magnified field of view highlighting the portion of area AM where each cell was identified for reconstruction. Right, reconstructions of single M2-projecting pyramidal neurons from area AM (magenta) receiving synaptic input from V1 axons (cyan); putative synaptic contacts are shown as blue circles. Number of putative synapses (markers with a distance below 0.25μm) noted above each reconstruction A-D: mouse #81234, E: mouse #81236, F-G: mouse #81233. Cells from deep layers: E and G. White scale bars = 50 μm. Black scale bars = 20 μm.

**Extended Data Figure 3-2.**
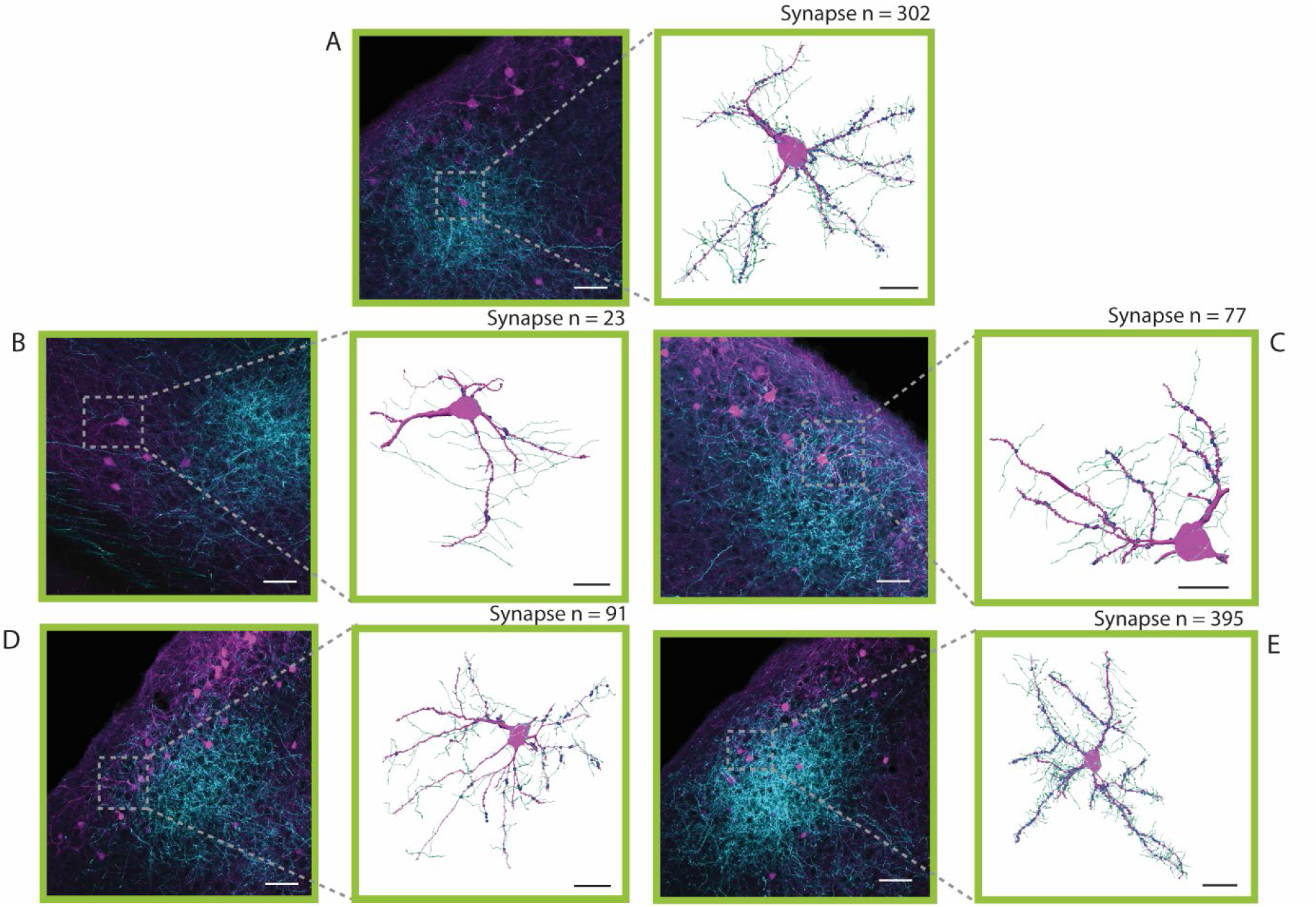
Five anatomical 3D reconstructions of V1 axons and M2-projecting neurons in extrastriate area AL. Left, inset, 20x magnification field of view highlighting the portion of area AL where the neurons were identified for reconstruction. Right, reconstructions of single M2-projecting pyramidal neurons from area AL (magenta) receiving synaptic input from V1 axons (cyan). Putative synaptic contacts are indicated by blue circles. The number of putative synapses (i.e. axon and dendrite segments separated by < 0.25μm) is noted above each reconstruction. A-C: mouse #81234, D-E: mouse #81236. B: deep layer neuron. White scale bars = 50 μm. Black scale bars = 20 μm.

**Extended Data Figure 3-3.**
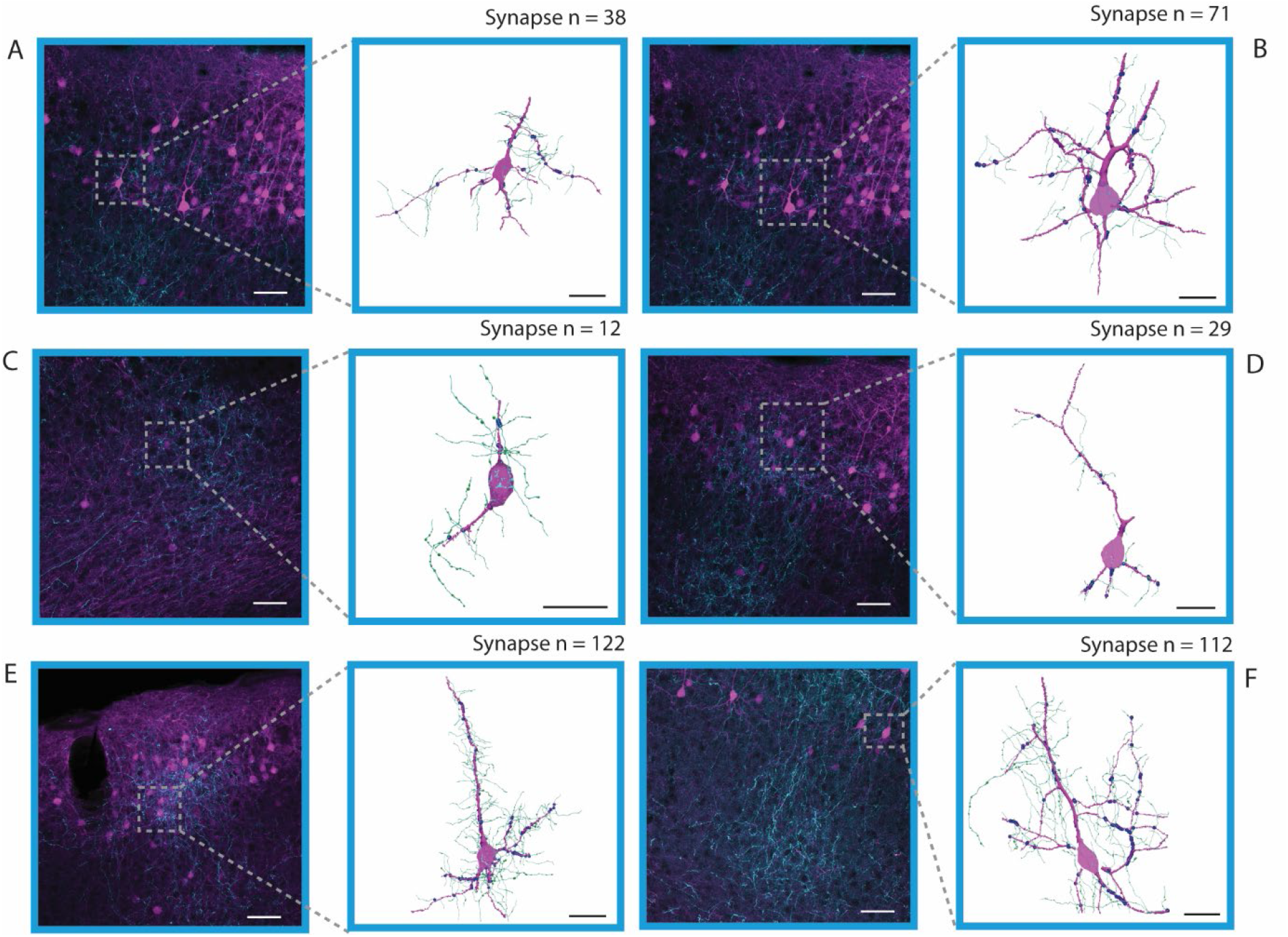
Six anatomical 3D reconstructions of M2-projecting neurons axons from V1 in extrastriate area PM. Left, inset, 20x magnified field of view highlighting the portion of area PM where each cell was identified for reconstruction. Right, reconstructions of single M2-projecting pyramidal neurons in area PM (magenta) receiving synaptic input from V1 axons (cyan); putative synaptic contacts are shown as blue circles. The number of putative synapses (markers with a distance < 0.25μm) is noted above each reconstruction. A-D: mouse #81234, E: mouse #81236, F: mouse #81233. Deep layer neurons: C. White scale bars = 50 μm. Black scale bars = 20 μm.

## References

Andermann, M. L., Kerlin, A. M., Roumis, D. K., Glickfeld, L. L., & Reid, R. C. (2011). Functional specialization of mouse higher visual cortical areas. Neuron, 72(6), 1025–1039. doi:10.1016/j.neuron.2011.11.013

Benevento, L. A., & Standage, G. P. (1982). Demonstration of lack of dorsal lateral geniculate nucleus input to extrastriate areas MT and visual 2 in the macaque monkey. Brain Res, 252(1), 161–166. doi:10.1016/0006-8993(82)90991-x

Brecht, M., Krauss, A., Muhammad, S., Sinai-Esfahani, L., Bellanca, S., & Margrie, T. W. (2004). Organization of rat vibrissa motor cortex and adjacent areas according to cytoarchitectonics, microstimulation, and intracellular stimulation of identified cells. J Comp Neurol, 479(4), 360–373. doi:10.1002/cne.20306

Burwell, R. D., & Amaral, D. G. (1998). Cortical afferents of the perirhinal, postrhinal, and entorhinal cortices of the rat. J Comp Neurol, 398(2), 179–205.

Callaway, E. M. (2004). Feedforward, feedback and inhibitory connections in primate visual cortex. Neural Netw, 17(5-6), 625–632. doi:10.1016/j.neunet.2004.04.004

Chafee, M. V., & Crowe, D. A. (2012). Thinking in spatial terms: decoupling spatial representation from sensorimotor control in monkey posterior parietal areas 7a and LIP. Front Integr Neurosci, 6, 112. doi:10.3389/fnint.2012.00112

Chen, L. L., Lin, L. H., Barnes, C. A., & McNaughton, B. L. (1994). Head-direction cells in the rat posterior cortex. ii. Contributions of visual and ideothetic information to the directional firing. Exp Brain Res, 101(1), 24–34.

Chen, X., Deangelis, G. C., & Angelaki, D. E. (2013). Diverse spatial reference frames of vestibular signals in parietal cortex. Neuron, 80(5), 1310–1321. doi:10.1016/j.neuron.2013.09.006

Coogan, T. A., & Burkhalter, A. (1993). Hierarchical organization of areas in rat visual cortex. J Neurosci, 13(9), 3749–3772.

Crowe, D. A., Averbeck, B. B., Chafee, M. V., & Georgopoulos, A. P. (2005). Dynamics of parietal neural activity during spatial cognitive processing. Neuron, 47(6), 885–891.

Donoghue, J. P., & Wise, S. P. (1982). The motor cortex of the rat: cytoarchitecture and microstimulation mapping. J Comp Neurol, 212(1), 76–88. doi:10.1002/cne.902120106

Douglas, R. J., & Martin, K. A. (2004). Neuronal circuits of the neocortex. Annu Rev Neurosci, 27, 419–451. doi:10.1146/annurev.neuro.27.070203.144152

Felleman, D. J., & Van Essen, D. C. (1991). Distributed hierarchical processing in the primate cerebral cortex. Cereb Cortex, 1(1), 1–47. doi:10.1093/cercor/1.1.1-a

Frost, D. O., & Caviness, V. S., Jr. (1980). Radial organization of thalamic projections to the neocortex in the mouse. J Comp Neurol, 194(2), 369–393. doi:10.1002/cne.901940206

Froudarakis, E., Fahey, P. G., Reimer, J., Smirnakis, S. M., Tehovnik, E. J., & Tolias, A. S. (2019). The Visual Cortex in Context. Annu Rev Vis Sci, 5, 317–339. doi:10.1146/annurev-vision-091517-034407

Garrett, M. E., Nauhaus, I., Marshel, J. H., & Callaway, E. M. (2014). Topography and areal organization of mouse visual cortex. J Neurosci, 34(37), 12587–12600. doi:10.1523/JNEUROSCI.1124-14.2014

Gilissen, S. R. J., Farrow, K., Bonin, V., & Arckens, L. (2021). Reconsidering the Border between the Visual and Posterior Parietal Cortex of Mice. Cereb Cortex, 31(3), 1675–1692. doi:10.1093/cercor/bhaa318

Glickfeld, L. L., Andermann, M. L., Bonin, V., & Reid, R. C. (2013). Cortico-cortical projections in mouse visual cortex are functionally target specific. Nat Neurosci, 16(2), 219–226. doi:10.1038/nn.3300

Goodale, M. A., & Milner, A. D. (1992). Separate visual pathways for perception and action. Trends in Neurosciences, 15(1), 20–25.

Han, X., Vermaercke, B., & Bonin, V. (2022). Diversity of spatiotemporal coding reveals specialized visual processing streams in the mouse cortex. Nat Commun, 13(1), 3249. doi:10.1038/s41467-022-29656-z

Harris, K. D., & Shepherd, G. M. (2015). The neocortical circuit: themes and variations. Nat Neurosci, 18(2), 170–181. doi:10.1038/nn.3917

Harvey, C. D., Coen, P., & Tank, D. W. (2012). Choice-specific sequences in parietal cortex during a virtual-navigation decision task. Nature, 484(7392), 62–68. doi:10.1038/nature10918

Hilgetag, C. C., Burns, G. A., O’Neill, M. A., Scannell, J. W., & Young, M. P. (2000). Anatomical connectivity defines the organization of clusters of cortical areas in the macaque monkey and the cat. Philos Trans R Soc Lond B Biol Sci, 355(1393), 91–110. doi:10.1098/rstb.2000.0551

Hilgetag, C. C., O’Neill, M. A., & Young, M. P. (1996). Indeterminate organization of the visual system. Science, 271(5250), 776–777. doi:10.1126/science.271.5250.776

Ho, J. W., Narduzzo, K. E., Outram, A., Tinsley, C. J., Henley, J. M., Warburton, E. C., & Brown, M. W. (2011). Contributions of area Te2 to rat recognition memory. Learn Mem, 18(7), 493–501. doi:10.1101/lm.2167511

Hovde, K., Gianatti, M., Witter, M. P., & Whitlock, J. R. (2018). Architecture and organization of mouse posterior parietal cortex relative to extrastriate areas. Eur J Neurosci. doi:10.1111/ejn.14280

Itokazu, T., Hasegawa, M., Kimura, R., Osaki, H., Albrecht, U. R., Sohya, K., Chakrabarti, S., Itoh, H., Ito, T., Sato, T. K., & Sato, T. R. (2018). Streamlined sensory motor communication through cortical reciprocal connectivity in a visually guided eye movement task. Nat Commun, 9(1), 338. doi:10.1038/s41467-017-02501-4

Keller, G. B., Bonhoeffer, T., & Hubener, M. (2012). Sensorimotor mismatch signals in primary visual cortex of the behaving mouse. Neuron, 74(5), 809–815. doi:10.1016/j.neuron.2012.03.040

Koganezawa, N., Gisetstad, R., Husby, E., Doan, T. P., & Witter, M. P. (2015). Excitatory Postrhinal Projections to Principal Cells in the Medial Entorhinal Cortex. J Neurosci, 35(48), 15860–15874. doi:10.1523/JNEUROSCI.0653-15.2015

Kolb, B., Buhrmann, K., McDonald, R., & Sutherland, R. J. (1994). Dissociation of the medial prefrontal, posterior parietal, and posterior temporal cortex for spatial navigation and recognition memory in the rat. Cereb Cortex, 4(6), 664–680.

Kolb, B., & Walkey, J. (1987). Behavioural and anatomical studies of the posterior parietal cortex in the rat. Behav Brain Res, 23(2), 127–145. doi:10166-4328(87)90050-7 [pii]

Krubitzer, L. A., & Seelke, A. M. (2012). Cortical evolution in mammals: the bane and beauty of phenotypic variability. Proc Natl Acad Sci U S A, 109 Suppl 1, 10647–10654. doi:10.1073/pnas.1201891109

Kaas, J. H., Krubitzer, L.A. (1991). The organization of extrastriate visual cortex. In B. Dreher, Robinson, S.R. (Ed.), Vision and Visual Dysfunction: Neuroanatomy of the Visual Pathways and Their Retinotopic Organization (Vol. 3, pp. 302–323). London: Nature Publishing Group.

Leinweber, M., Ward, D. R., Sobczak, J. M., Attinger, A., & Keller, G. B. (2017). A Sensorimotor Circuit in Mouse Cortex for Visual Flow Predictions. Neuron, 95(6), 1420–1432 e1425. doi:10.1016/j.neuron.2017.08.036

Marshel, J. H., Garrett, M. E., Nauhaus, I., & Callaway, E. M. (2011). Functional specialization of seven mouse visual cortical areas. Neuron, 72(6), 1040–1054. doi:10.1016/j.neuron.2011.12.004

Maunsell, J. H., & Newsome, W. T. (1987). Visual processing in monkey extrastriate cortex. Annu Rev Neurosci, 10, 363–401. doi:10.1146/annurev.ne.10.030187.002051

McNaughton, B. L., Leonard, B., & Chen, L. (1989). Cortical-hippocampal interactions and cognitive mapping: A hypothesis based on reintegration of the parietal and inferotemporal pathways for visual processing. Psychobiology, 17(3), 230–235. doi:10.1007/BF03337774

Miller, M. W., & Vogt, B. A. (1984). Direct connections of rat visual cortex with sensory, motor, and association cortices. J Comp Neurol, 226(2), 184–202. doi:10.1002/cne.902260204 [doi]

Mlinar, E. J., & Goodale, M. A. (1984). Cortical and tectal control of visual orientation in the gerbil: evidence for parallel channels. Exp Brain Res, 55(1), 33–48. doi:10.1007/BF00240496

Montero, V. M. (1993). Retinotopy of cortical connections between the striate cortex and extrastriate visual areas in the rat. Exp Brain Res, 94(1), 1–15.

Montero, V. M., & Jian, S. (1995). Induction of c-fos protein by patterned visual stimulation in central visual pathways of the rat. Brain Res, 690(2), 189–199. doi:10.1016/0006-8993(95)00620-6

Nassi, J. J., & Callaway, E. M. (2009). Parallel processing strategies of the primate visual system. Nat Rev Neurosci, 10(5), 360–372. doi:10.1038/nrn2619

Niell, C. M. (2015). Cell types, circuits, and receptive fields in the mouse visual cortex. Annu Rev Neurosci, 38, 413–431. doi:10.1146/annurev-neuro-071714-033807

Nitz, D. A. (2006). Tracking route progression in the posterior parietal cortex. Neuron, 49(5), 747–756.

Olavarria, J., Mignano, L. R., & Van Sluyters, R. C. (1982). Pattern of extrastriate visual areas connecting reciprocally with striate cortex in the mouse. Exp Neurol, 78(3), 775–779. doi:10.1016/0014-4886(82)90090-5

Olavarria, J., & Montero, V. M. (1989). Organization of visual cortex in the mouse revealed by correlating callosal and striate-extrastriate connections. Vis Neurosci, 3(1), 59–69. doi:10.1017/s0952523800012517

Olsen, G. M., Hovde, K., Kondo, H., Sakshaug, T., Somme, H. H., Whitlock, J. R., & Witter, M. P. (2019). Organization of Posterior Parietal-Frontal Connections in the Rat. Front Syst Neurosci, 13, 38. doi:10.3389/fnsys.2019.00038

Paxinos, G., Franklin K. (2012). Paxino’s and Franklin’s the Mouse Brain in Stereotaxic Coordinates (4th ed.): Academic Press.

Reep, R. L., Goodwin, G. S., & Corwin, J. V. (1990). Topographic organization in the corticocortical connections of medial agranular cortex in rats. J Comp Neurol, 294(2), 262–280. doi:10.1002/cne.902940210

Rosa, M. G., & Krubitzer, L. A. (1999). The evolution of visual cortex: where is V2? Trends Neurosci, 22(6), 242–248. doi:10.1016/s0166-2236(99)01398-3

Rozzi, S., Ferrari, P. F., Bonini, L., Rizzolatti, G., & Fogassi, L. (2008). Functional organization of inferior parietal lobule convexity in the macaque monkey: electrophysiological characterization of motor, sensory and mirror responses and their correlation with cytoarchitectonic areas. Eur J Neurosci, 28(8), 1569–1588. doi:10.1111/j.1460-9568.2008.06395.x

Salin, P. A., & Bullier, J. (1995). Corticocortical connections in the visual system: structure and function. Physiol Rev, 75(1), 107–154. doi:10.1152/physrev.1995.75.1.107

Save, E., Poucet, B., Foreman, N., & Buhot, M. C. (1992). Object exploration and reactions to spatial and nonspatial changes in hooded rats following damage to parietal cortex or hippocampal formation. Behav Neurosci, 106(3), 447–456.

Schneider, G. E. (1969). Two visual systems. Science, 163(3870), 895–902. doi:10.1126/science.163.3870.895

Simmons, P. A., Lemmon, V., & Pearlman, A. L. (1982). Afferent and efferent connections of the striate and extrastriate visual cortex of the normal and reeler mouse. J Comp Neurol, 211(3), 295–308. doi:10.1002/cne.902110308

Sinnamon, H. M., & Galer, B. S. (1984). Head movements elicited by electrical stimulation of the anteromedial cortex of the rat. Physiol Behav, 33(2), 185–190. doi:10.1016/0031-9384(84)90098-2

Tees, R. C. (1999). The effects of posterior parietal and posterior temporal cortical lesions on multimodal spatial and nonspatial competencies in rats. Behav Brain Res, 106(1-2), 55–73. doi:10.1016/s0166-4328(99)00092-3

Tervo, D. G., Hwang, B. Y., Viswanathan, S., Gaj, T., Lavzin, M., Ritola, K. D., Lindo, S., Michael, S., Kuleshova, E., Ojala, D., Huang, C. C., Gerfen, C. R., Schiller, J., Dudman, J. T., Hantman, A. W., Looger, L. L., Schaffer, D. V., & Karpova, A. Y. (2016). A Designer AAV Variant Permits Efficient Retrograde Access to Projection Neurons. Neuron, 92(2), 372–382. doi:10.1016/j.neuron.2016.09.021

Tombaz, T., Dunn, B. A., Hovde, K., Cubero, R. J., Mimica, B., Mamidanna, P., Roudi, Y., & Whitlock, J. R. (2020). Action representation in the mouse parieto-frontal network. Sci Rep, 10(1), 5559. doi:10.1038/s41598-020-62089-6

van Groen, T., & Wyss, J. M. (1992). Connections of the retrosplenial dysgranular cortex in the rat. J Comp Neurol, 315(2), 200–216.

Viaro, R., Maggiolini, E., Farina, E., Canto, R., Iriki, A., D’Ausilio, A., & Fadiga, L. (2021). Neurons of rat motor cortex become active during both grasping execution and grasping observation. Curr Biol, 31(19), 4405–4412 e4404. doi:10.1016/j.cub.2021.07.054

Vogt, B. A., & Miller, M. W. (1983). Cortical connections between rat cingulate cortex and visual, motor, and postsubicular cortices. J Comp Neurol, 216(2), 192–210. doi:10.1002/cne.902160207

Wang, Q., & Burkhalter, A. (2007). Area map of mouse visual cortex. J Comp Neurol, 502(3), 339–357. doi:10.1002/cne.21286

Wang, Q., Gao, E., & Burkhalter, A. (2011). Gateways of ventral and dorsal streams in mouse visual cortex. J Neurosci, 31(5), 1905–1918. doi:10.1523/JNEUROSCI.3488-10.2011

Wang, Q., Sporns, O., & Burkhalter, A. (2012). Network analysis of corticocortical connections reveals ventral and dorsal processing streams in mouse visual cortex. J Neurosci, 32(13), 4386–4399. doi:10.1523/JNEUROSCI.6063-11.2012

Whitlock, J. R., Pfuhl, G., Dagslott, N., Moser, M. B., & Moser, E. I. (2012). Functional split between parietal and entorhinal cortices in the rat. Neuron, 73(4), 789–802. doi:10.1016/j.neuron.2011.12.028S0896-6273(12)00078-5 [pii]

Wilber, A. A., Clark, B. J., Forster, T. C., Tatsuno, M., & McNaughton, B. L. (2014). Interaction of egocentric and world-centered reference frames in the rat posterior parietal cortex. J Neurosci, 34(16), 5431–5446. doi:10.1523/JNEUROSCI.0511-14.2014

Wouterlood, F. G., Boekel, A. J., Kajiwara, R., & Belien, J. A. (2008). Counting contacts between neurons in 3D in confocal laser scanning images. J Neurosci Methods, 171(2), 296–308. doi:10.1016/j.jneumeth.2008.03.014

Yang, F. C., Jacobson, T. K., & Burwell, R. D. (2017). Single neuron activity and theta modulation in the posterior parietal cortex in a visuospatial attention task. Hippocampus, 27(3), 263–273. doi:10.1002/hipo.22691

Zhang, S., Xu, M., Kamigaki, T., Hoang Do, J. P., Chang, W. C., Jenvay, S., Miyamichi, K., Luo, L., & Dan, Y. (2014). Selective attention. Long-range and local circuits for top-down modulation of visual cortex processing. Science, 345(6197), 660–665. doi:10.1126/science.1254126

Zingg, B., Hintiryan, H., Gou, L., Song, M. Y., Bay, M., Bienkowski, M. S., Foster, N. N., Yamashita, S., Bowman, I., Toga, A. W., & Dong, H. W. (2014). Neural networks of the mouse neocortex. Cell, 156(5), 1096–1111. doi:10.1016/j.cell.2014.02.023

